# Establishing Rod-Shape from Spherical, Peptidoglycan-Deficient Bacterial Spores

**DOI:** 10.1101/818641

**Authors:** Huan Zhang, Garrett A. Mulholland, Sofiene Seef, Shiwei Zhu, Jun Liu, Tâm Mignot, Beiyan Nan

**Affiliations:** Department of Biology, Texas A&M University, College Station, TX 77843, USA; Laboratoire de Chimie Bactérienne, CNRS - Université Aix Marseille UMR7283, Institut de Microbiologie de la Méditerranée, 13009 Marseille, France; Department of Microbial Pathogenesis, Yale University School of Medicine, New Haven, CT 06536, USA; Microbial Sciences Institute, Yale University, West Haven, CT 06516, USA

**Author notes:** Corresponding author: Beiyan Nan, 306C BSBE, 3258 TAMU, College Station, TX77843.

**Keywords:** germination, Rod system, MreB, GTPase, gliding motor

## Abstract

Chemical-induced spores of the Gram-negative bacterium *Myxococcus xanthus* are peptidoglycan (PG)-deficient. It is unclear how these spherical spores germinate into rod-shaped, walled cells without preexisting PG templates. We found that germinating spores first synthesize PG randomly on spherical surfaces. MglB, a GTPase-activating protein, forms a cluster that surveys the status of PG growth and stabilizes at one future cell pole. Following MglB, the Ras family GTPase MglA localizes to the second pole. MglA directs molecular motors to transport the bacterial actin homolog MreB and the Rod PG synthesis complexes away from poles. The Rod system establishes rod-shape by elongating PG at nonpolar regions. Thus, the interaction between GTPase, cytoskeletons and molecular motors provides a mechanism for the *de novo* establishment of rod-shape in bacteria.

**Significance:** Spheres and rods are among the most common shapes adopted by walled bacteria, in which the peptidoglycan (PG) cell wall largely determines cell shape. When induced by chemicals, rod-shaped vegetative cells of the Gram-negative bacterium *Myxococcus xanthus* thoroughly degrade their PG and shrink into spherical spores. As these spores germinate, rod-shaped cells are rebuilt without preexisting templates, which provides a rare opportunity to visualize *de novo* PG synthesis and bacterial morphogenesis. In this study, we investigated how spherical spores germinate into rods and elucidated a system for rod-shape morphogenesis that includes the Rod PG synthesis system, a GTPase-GAP pair, the MreB cytoskeleton and a molecular motor.

---

Morphogenesis is a fundamental feature of cells. Compared to spheres that are symmetric in all directions, rods are asymmetric and polarized. For most rod-shaped bacteria, the peptidoglycan (PG) cell wall defines cell geometry, which is assembled by two major enzymatic systems. The Rod system consists of RodA, a SEDS-family PG polymerase, PBP2, a member of the class B penicillin-binding proteins (bPBPs), and MreB, a bacterial actin homolog that orchestrates the activities of the Rod complexes in response to local cell curvature (1). In contrast, class A PBPs (aPBPs) contribute to PG growth independent of MreB (2, 3).

*Myxococcus xanthus*, a rod-shaped Gram-negative bacterium, utilizes polarized geometry for directed locomotion. MglA, a Ras family small GTPase, controls the direction of gliding motility (4–7). As cells move, GTP-bound MglA forms large clusters at leading cell poles, whereas GDP-bound MglA distributes homogeneously in the cytoplasm (4, 6, 7). MglA-GTP stimulates the assembly of the gliding machineries through direct interaction with MreB (7–10) and directs the gliding machineries toward lagging cell poles (5). Consequently, the gliding machineries carry MreB filaments as they move rapidly in the membrane (11–13). The activity of MglA is regulated by its cognate GTPase-activating protein (GAP), MglB, which forms large clusters at lagging cell poles. MglB activates the GTPase activity of MglA, expelling MglA-GTP, and thus the assembled gliding machineries, from lagging poles (4, 6). Overall, polarized localization and activities of MglA and MglB ensure that the of gliding machineries generate propulsion by moving from poles to nonpolar regions (5, 7, 14).

Some rod-shaped bacteria change their geometry through sporulation. In Firmicutes such as Bacilli and Clostridia, the morphological differentiation from rod-shaped vegetative cells to oval spores begins with an asymmetric division, resulting in the formation of a smaller endospore wholly contained within a larger mother cell. In contrast to endospore-forming bacteria, *M. xanthus* produces spores using two division-independent mechanisms. First, groups of vegetative cells can aggregate on solid surfaces and build spore-filled fruiting bodies in a few days (15). Second, individual *M. xanthus* cells can form dispersed, spherical spores within hours in response to chemical signals, such as glycerol (16). Unlike endospores that contain intact and often thickened PG (17, 18), PG is thoroughly degraded in glycerol-induced *M. xanthus* spores (19). Without the polarity defined by PG, the mechanism by which these spores elongate into rods remains largely unknown.

## Results

### Two-phase morphological transition during *M. xanthus* spore germination

Overnight induction of wild-type cells by 1 M glycerol produced sonication-resistant spores with length to width aspect ratios (L/W) of 1.56 ± 0.36 (n = 789), among which 40.9% are approximately spherical (L/W ≤ 1.3). Overall, the L/W values of most (85.4%) spores were lower than 2. As spores germinated, the morphological transition progressed in a two-phase manner. In the first hour (Phase I), L/W did not change significantly (*p* = 0.57, Fig. 1A, 1B, *SI Appendix*, Movie S1, Table S1). After 1 h, L/W increased sharply as emerging cells transformed into rods (Phase II). Importantly, 40.2% (n = 244) spores did not initiate elongation along their original long axes (Fig. 1C, *SI Appendix,* Movie S2), indicating that although not perfectly spherical, the geometry of mature spores does not predetermine the polarity of emerging cells. 70.2% (n = 198) of emerging cells reached the dimensions of vegetative cells by 3 h. After 8 h, the whole population of emerging cells is indistinguishable from vegetative cells (L/W = 5.55 ± 1.12, n = 233. Fig. 1A, 1B, *SI Appendix,* Movie S1, Table S1).

**Fig. 1.**
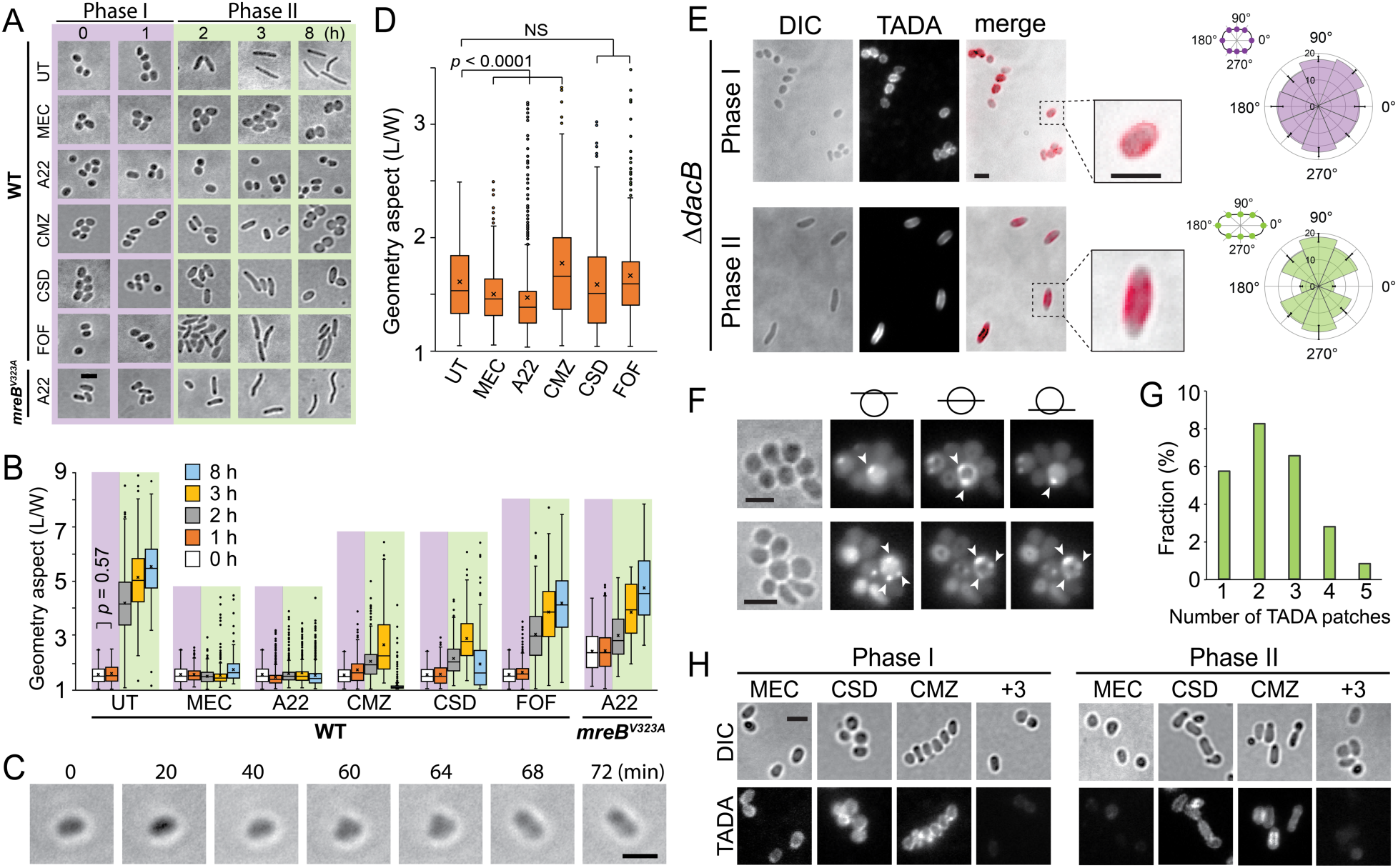
PG polymerization by the Rod system is essential for *de novo* establishment of rod-shape. **A)** Morphological changes of untreated (UT) and inhibitor-treated spores in the germination process. A22 (10 μg/ml), mecillinam (MEC, 100 μg/ml), cefmetazole (CMZ, 5 mg/ml), cefsulodin (CSD, 5 mg/ml) and fosfomycin (FOF, 8 mg/ml). **B)** Quantitative analysis of the germination progress using the aspect ratios (L/W) of spores/cells. Boxes indicate the 25^th^ - 75^th^ percentiles, whiskers the 5^th^ - 95^th^ percentiles. In each box, the midline indicates the median and × indicates the mean (same below, Table S1). Outlier data points are shown as individual dots above and below the whiskers. **C)** An oval spore initiates elongation along its short axes (Movie S2). **D)** Phase I spores become more spherical after 1-h treatments by A22 and mecillinam. Cefmetazole-treated spores initiate elongation earlier than untreated ones. **E)** Patterns of PG growth in both phases of germination were visualized by TADA in *ΔdacB* spores. The average and standard deviation of TADA intensity were calculated from 20 spores/cells in the diagrams to the right (same below). **F)** Imaged at different focal planes, 22.0% of Phase I spores show bright TADA patches (arrows) that position randomly on spore surfaces. **G)** Among these 22.0% spores, many contain multiple TADA patches. **H)** Compared to untreated (UT) *ΔdacB* spores, while neither MEC, CMZ or CSD is able to block PG growth, the combination of all three antibiotics (+3) abolishes PG growth in Phase I of germination. In contrast, MEC alone is sufficient to inhibit PG growth in Phase II. Scale bars, 2 μm. *p* values were calculated using the Student paired *t* test with a two-tailed distribution (same below). NS, nonsignificant difference.

Using cryo-electron tomography (cryo-ET), we confirmed that mature glycerol-induced spores do not retain PG (*SI Appendix,* Fig. S1). To investigate the role of PG growth in germination, we treated spores with several inhibitors for PG assembly. In the presence of A22, an inhibitor of MreB in the Rod system, spores failed to germinate into rods as their L/W ratios did not increase within 8 h. In contrast, when the wild-type MreB was replaced by MreB^V323A^, an A22-resistant variant (9), spores were able to elongate into rods in the presence of A22, indicating that A22 inhibits germination specifically through MreB (Fig. 1A, 1B, *SI Appendix,* Table S1). Similarly, mecillinam that inhibits PBP2 in the Rod system also blocked germination (Fig. 1A, 1B, *SI Appendix,* Table S1). Both A22 and mecillinam-treated spores were viable because they were able to grow into rods when transferred into inhibitor-free medium. Since these spores became even more spherical in Phase I (Fig. 1D), neither A22 nor mecillinam blocked the hydrolysis of spore coats that maintain the shape of oval spores. In the presence of cefsulodin that inhibits PBP1A/B, and cefmetazole that inhibits all PBPs except PBP2, spores were able to form rods, albeit the elongation rate was slower (Fig. 1A, 1B, *SI Appendix,* Table S1). Although not essential for the initiation of cell elongation, aPBPs still contribute to the maintenance of rod shape. Despite successful elongation in early Phase II (1 – 3 h), 57.2% (n = 215) cefsulodin-treated and 96.6% (n = 203) cefmetazole-treated emerging cells retrogressed to spheres after 8-h treatments (Fig. 1A, 1B, *SI Appendix,* Table S1), which is in agreement with a recent report that aPBPs are required for the mechanical stability of cells (20). L/W values of the cefmetazole-treated spores increased significantly in Phase I of germination (*p* < 0.0001, Fig. 1B, 1D, *SI Appendix,* Table S1), suggesting that cells elongate earlier when PBP2 is dominant over other PBPs. Vegetative *M. xanthus* cells are sensitive to fosfomycin, an antibiotic that inhibits the production of UDP-MurNAc, a precursor of PG (11). As spores preserve PG precursors (19), they were able to elongate into rods in the presence of fosfomycin, albeit at reduced rates (Fig. 1A, 1B, *SI Appendix,* Table S1). Taken together, PG polymerization by the Rod system is essential for the establishment of rod-shape.

Bacterial cells incorporate single D-amino acid-based fluorescent probes into PG using D, D- and L, D-transpeptidases that catalyze the last steps of PG polymerization (21). Such fluorescent D-amino acids could serve as proxy reporters for PG growth. We next visualized the patterns of PG growth using a fluorescent D-amino acid, TAMRA 3-amino-D-alanine (TADA) (22), to label newly synthesized PG. To enhance labeling efficiency, we deleted the *dacB* gene (*mxan_3130*), which encodes a D-Ala-D-Ala carboxypeptidase (23). The resulted *ΔdacB* cells showed identical morphology to the wild-type ones and produced sonication-resistant spores. The *ΔdacB* spores showed minor delay in germination and efficient TADA incorporation (Fig. 1E, *SI Appendix,* Fig. S2 and Table S1). Although L/W of spores did not change in Phase I, PG had started to grow. The surfaces of most Phase I spores (78.0%, n = 600) were evenly labeled by TADA (Fig. 1E). The remaining 22.0% of spores showed bright patches of TADA on their surfaces (Fig. 1F). However, these patches do not likely register future poles because 47.0% (n = 132) of such spores contained more than two TADA patches and these patches positioned randomly on spore surfaces (Fig. 1F, 1G). In contrast, as cells grew into rods, TADA was incorporated heavily at nonpolar regions and fluorescence signals were generally absent at cell poles (Fig. 1E). The patterns of PG growth indicate that spores first synthesize PG on their spherical surfaces in Phase I and then break symmetry in Phase II by growing PG at nonpolar regions.

Neither mecillinam, cefsulodin or cefmetazole was able to block TADA incorporation in Phase I of germination. However, a treatment by all three antibiotics abolished TADA incorporation (Fig. 1H), indicating that both aPBPs and the Rod system contribute to the isotropic PG growth in Phase I. In contrast, mecillinam, but not cefsulodin or cefmetazole, blocked TADA incorporation in Phase II of germination (Fig. 1H). The coordination between aPBPs and the Rod system still remains to be fully understood. On the one hand, since cefsulodin and cefmetazole reduce the elongation rates of emerging cells (Fig. 1A, 1B, *SI Appendix,* Table S1), aPBPs still participate in PG synthesis in Phase II of germination. But on the other hand, the activities of aPBPs might depend on the Rod system because mecillinam alone is sufficient to block TADA incorporation in Phase II. Consistent with a recent report that cells reduce their diameter when the Rod system becomes dominant over aPBPs (24), emerging cells continued to grow in length but shrink in width in Phase II (Fig. S3, *SI Appendix,* Table S1). These results confirm that while both aPBPs and the Rod system participate PG synthesis in germination, the Rod system plays major roles in cell elongation.

### MglA and MglB are required for rapid cell elongation

To investigate how *M. xanthus* spores establish rod shape *de novo*, we tested the potential roles of polar-localized motility regulators. *ΔmglA* and *ΔmglB* cells were able to form sonication-resistant spores but their spores showed severe delays in elongation, especially in early Phase II of germination. After 3 h, only 15.7% of the *ΔmglA* (n= 140) and 10.4% of *ΔmglB* (n = 298) cells reached the vegetative aspect ratio (comparing to 70.2% of wild-type cells, Fig. 2A, 2B, *SI Appendix,* Table S1). In contrast, deleting *romR* and *plpA*, the genes encode two additional polar-localized motility regulators (25–27), only caused minor delay in germination (*SI Appendix,* Fig. S2 and Table S1). Both the *ΔmglA* and *ΔmglB* spores were able to elongate in length and shrink in width, albeit at significantly lower rates (Fig. 2A, 2B, *SI Appendix,* Table S1, Fig. S3), indicating that PG growth by the Rod complex still occurred. Strikingly different from wild-type spores that maintained relatively smooth cell surfaces during germination, cells from the *ΔmglA* and *ΔmglB* spores showed pronounced bulges at nonpolar regions in early Phase II, appearing to have multiple cell poles (Fig. 2A, 2C, *SI Appendix,* Movie S3). However, this morphological defect was largely corrected after prolonged growth (8 h) (Fig. 2A), implying that a system independent of MglA and MglB was able to generate rod shape, although much less robustly. To determine how MglA and MglB regulate germination, we investigated the spores that expressed the MglA^Q82L^ variant as the sole source of MglA, under the control of the native promoter of the *mglBA* operon. MglA^Q82L^ expresses normally but is unable to hydrolyze GTP (6). Spores expressing wild-type MglB and MglA^Q82L^ showed both a severe delay in cell elongation and bulged surfaces on emerging cells, similar to the *ΔmglA* and *ΔmglB* spores (Fig. 2A, 2B, *SI Appendix,* Table S1). Surprisingly, overproducing MglB (*mglB^OE^*), which potentially overstimulates the GTPase activity of MglA, caused similar defects during germination (Fig. 2A, 2B, *SI Appendix,* Table S1). Thus, fine-tuned GTPase activity of MglA is required for rapid cell elongation and MglB functions through MglA.

**Fig. 2.**
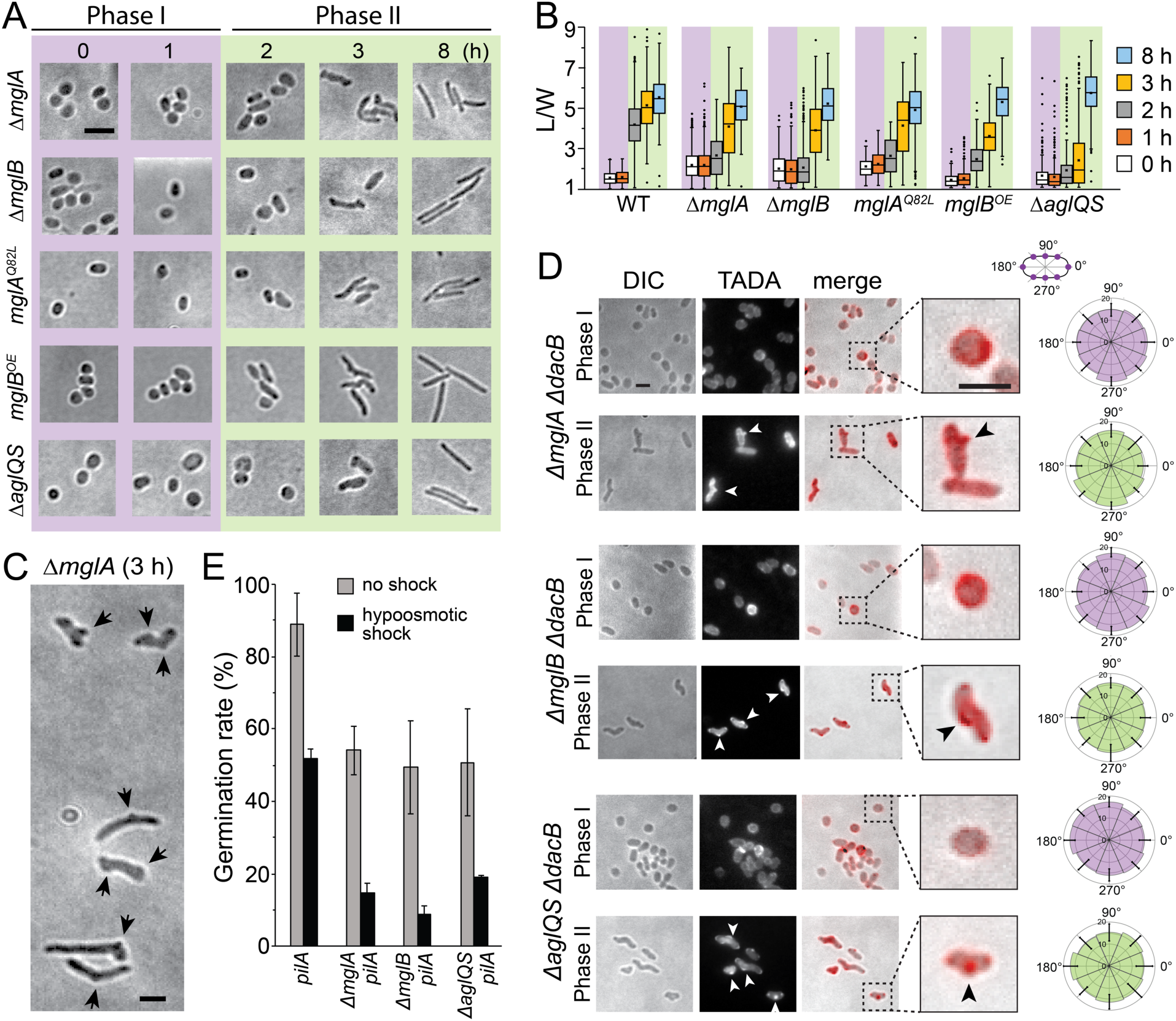
MglA and MglB are required for rapid cell elongation. **A)** Emerging cells from *ΔmglA*, *ΔmglB, mglA^Q82L^* and *mglB^OE^* spores and *ΔaglQS* pseudospores show significant delay in elongation and bulges on cell surfaces in Phase II of germination. **B)** Quantitative analysis of the germination progress. **C)** A representative image of the altered morphology of the emerging *ΔmglA* cells after 3-h of germination. Arrows point to the bulges on cell surfaces. **D)** The disruption of either the MglA-MglB polar axis (*ΔmglA ΔdacB* and *ΔmglB ΔdacB*) or the gliding motor (*ΔaglQS ΔdacB*) resulted in significantly stronger PG growth at cell poles and budges (arrows) in Phase II. Quantitative analysis of TADA fluorescence is shown on the right. **E)** The disruption of either the MglA-MglB polar axis (*ΔmglA pilA::tet* and *ΔmglB pilA::tet*) or the gliding motor (*ΔaglQS pilA::tet*) significantly decreases the survival rate of germinating spores (gray bars), especially under osmotic stress (black bars). Scale bars, 2 μm.

Both the delayed morphological transition and bulged surfaces of the emerging *ΔmglA* and *ΔmglB* cells suggest that MglA and MglB might regulate PG growth during germination. *ΔmglA ΔdacB* and *ΔmglB ΔdacB* spores were able to grow PG in an isotropic manner in Phase I, indistinguishable from the *ΔdacB* spores (Fig. 2D). However, emerging cells from both mutant spores displayed elevated PG growth at cell poles and bulges in Phase II (Fig. 2D).

Delayed elongation and uneven PG growth during germination could reduce the viability of *ΔmglA* and *ΔmglB* spores, especially under osmotic stresses. To test this hypothesis, we enumerated spores in cell-counting chambers and calculated their viability by dilution plating. To simplify cell-counting, *pilA* was disrupted by plasmid insertion (*pilA::tet*) to reduce cell aggregation caused by exopolysaccharides production (28). As shown in Fig. 2E, compared to the *pilA::tet* spores that 88.9 ± 8.7% (calculated from 3 independent experiments, same below) formed colonies, both *ΔmglA pilA::tet and ΔmglB pilA::tet* spores formed colonies at reduced rates (54.1 ± 6.7 and 49.4 ± 12.9%, respectively). To further test if altered growth pattern reduces the strength of PG, we allow spores to germinate for 1 h, then incubated emerging cells in a hypoosmotic buffer (20 mM Tris-HCl, pH 7.6) for 1 h before plating. Hypoosmotic shock reduced the survival rate of *pilA::tet* spores to 51.8 ± 2.6%, indicating that the still-growing PG in germination Phase II is sensitive to osmotic stress. Strikingly, after hypoosmotic shock, less than 15% of *ΔmglA pilA::tet* (14.5 ± 2.8%) and *ΔmglB pilA::tet* (8.7 ± 2.4%) spores formed colonies (Fig. 2E). Taken together, the MglA-MglB polarity axis regulates PG growth in Phase II of germination, which plays important roles in the survival of glycerol-induced *M. xanthus* spores.

### MglB stabilizes at the first future pole

We expressed YFP-labeled MglA as merodiploids in the wild-type background (6, 9) and mCherry-labeled MglB (stably expressed, *SI Appendix,* Table S1, Fig. S2) in the *ΔmglB* mutant and correlated their localization patterns with germination progress (L/W). Spores from neither strain showed significant defects in germination (*SI Appendix,* Table S1, Fig. S2). 94.1% (n = 152) of Phase I spores (L/W ≤ 2) contained one or two MglB clusters (Fig. 3A, 3B). In phase II spores (L/W > 2), this ratio increased to 100% (n = 120). In contrast, MglA did not form clusters until Phase II, when 54.2% of emerging cells contained one or two MglA clusters (Fig. 3A, 3B). Thus, during germination, MglB establishes polarized localization before MglA.

**Fig. 3.**
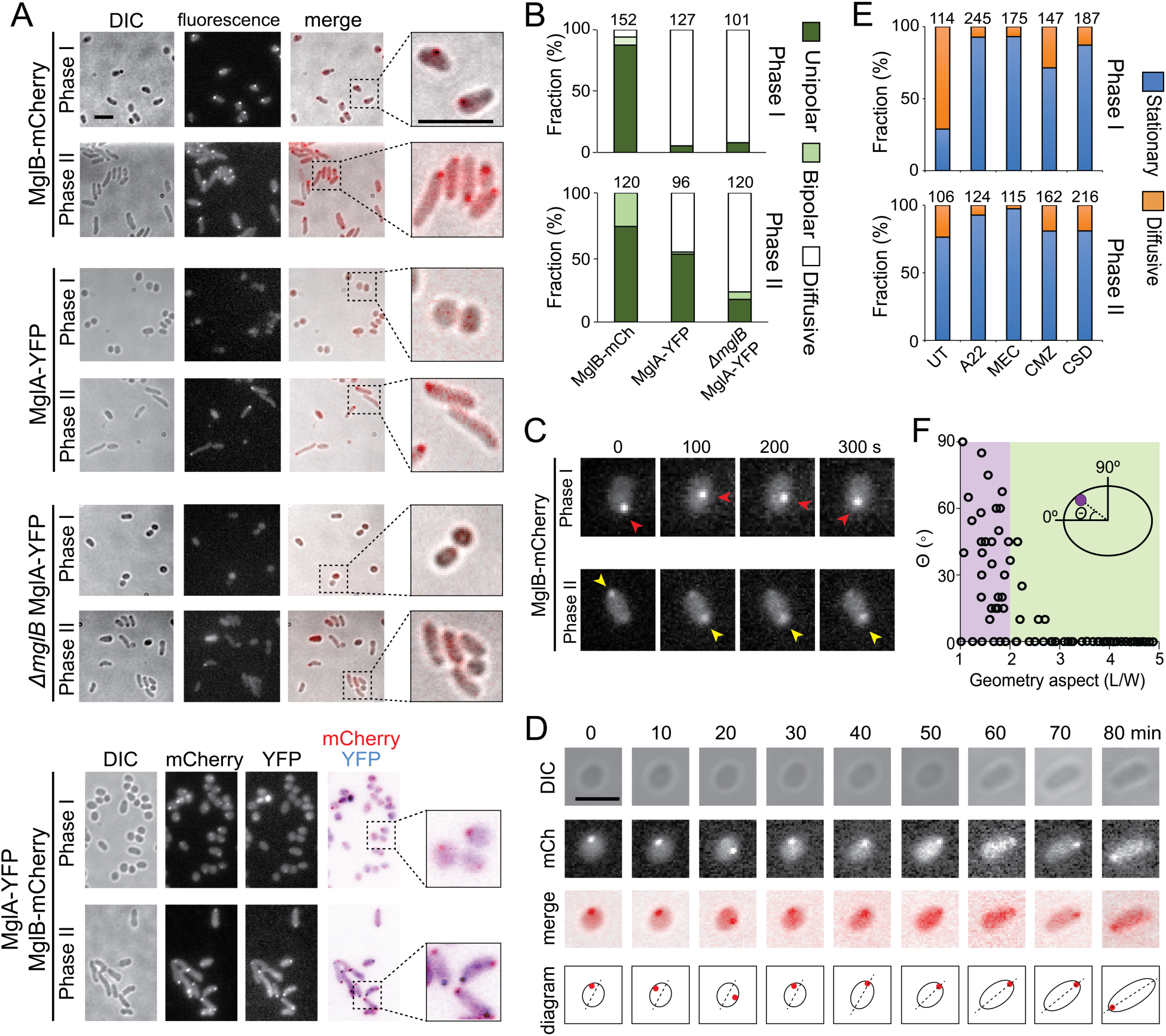
MglB stabilizes at the first future pole. **A)** While MglB-mCherry forms clusters in germination Phase I, MglA-YFP starts to forms clusters in Phase II. In the absence of MglB, MglA-YFP forms significantly fewer clusters. MglA and MglB clusters stabilize at opposite cell poles. **B)** Quantification of the sequential stabilization of MglB and MglA clusters. The total number of spores/cells analyzed for each strain is shown on top of each bar. **C)** The “wandering” dynamics of MglB clusters (red arrow) in Phase I spores (Movie S4). In Phase II, MglB clusters (yellow arrow) stabilized at cell poles and oscillated between opposite poles (Movie S6). **D)** As the MglB cluster stabilizes at one future pole, the emerging cell starts to elongate (Movie S5). **E)** Inhibitors of PG synthesis, A22, MEC, CMZ and CSD, all inhibit the wandering of MglB in Phase I. In contrast, only A22 and MEC, the inhibitors of the Rod system, inhibit the wandering of MglB significantly in Phase II. For each treatment, the total number of MglB clusters analyzed is shown on top of each bar. UT, untreated. **F)** The stabilization of MglB clusters does not depend on local curvature. Scales bars, 2 μm.

To test if the clusters of MglB in Phase I spores mark the polarity inherited from previous vegetative cells, we imaged MglB clusters at 0.05 Hz. Surprisingly, the majority of MglB clusters in Phase I spores was highly dynamic (Fig. 3C, *SI Appendix,* Movie S4). Among 114 MglB clusters in Phase I spores, 22.9% remained stationary, and 77.1% showed typical diffusion, with diffusion coefficients (*D_MglB_*) of 1.05 × 10^-4^ ± 4.62 × 10^-5^ μm^2^/s. These “wandering” MglB clusters were observed in both the approximately spherical (L/W < 1.3) and oval spores (1.3 < L/W ≤ 2), which supports our hypothesis that, regardless of their geometry, polarity is not yet established in Phase I spores.

As germination progressed, MglB clusters started to stabilize. Strikingly, once stabilized, the locations of MglB clusters became one of the cell poles for cell elongation (Fig. 3D, *SI Appendix,* Movie. S5). In Phase II of germination (L/W > 2), the population of stationary MglB clusters increased from 22.9% to 76.4% (n = 106, Fig. 3E). Stabilized MglB clusters began to oscillate between newly established poles (Fig. 3C, *SI Appendix,* Movie S6), which might provide a mechanism to ensure that MglB occupies each future cell pole for an equal amount of time. As MglB clusters stabilized, MglA started to form clusters. The formation of MglA-YFP clusters was delayed significantly in the *ΔmglB* background, and only 23.3% (n = 120) of emerging cells contained MglA clusters in Phase II of germination (Fig. 3A, 3B). Among the 204 cells in which both proteins formed single clusters, 176 (86.3%) positioned MglA and MglB clusters at opposite poles (Fig. 3A, 3B). Thus, spherical spores start to elongate into rods along the axes established by the sequential stabilization of MglB and MglA clusters.

To investigate whether the stabilization of MglB clusters is predetermined by local cell curvatures, we quantified the localization of stationary MglB clusters with regard to the geometry of spores. We divide each spore/cell envelope into four quarters. In the quarter that contained stationary MglB clusters, we defined the long and short axes as 0° and 90°, which mark the local curvature that shows the highest and lowest similarity to the poles of vegetative cells, respectively. As shown in Fig. 3F, MglB clusters stabilized randomly in Phase I spores, indicating that local curvature does not dictate the localization of MglB. After the stabilization of MglB, the sites harboring MglB clusters transformed into cell poles (0°) in Phase II (Fig. 3F).

MglB clusters could stabilize at the sites where PG synthesis has completed or not yet initiated. We ruled out the second possibility because the majority of MglB clusters (76.4%, n = 106) stabilizes at poles in Phase II (Fig. 3A, 3B), where almost no PG growth was observed later (Fig. 1E). In Phase I of germination, the population of static MglB clusters increased dramatically in the presence of A22, mecillinam, cefmetazole and cefsulodin (Fig. 3E), indicating that active PG growth prevents MglB clusters from settling down. Consistent with our finding that the Rod system becomes the dominant system for PG growth in Phase II (Fig. 1H), A22 and mecillinam further reduced the small population of diffusive MglB clusters in early Phase II, while cefmetazole and cefsulodin did not show significant effects (Fig. 3E). Taken together, it is the progress of PG growth, rather than the geometry of the spore, that regulates the dynamic of MglB clusters. As MglB clusters only stabilize at the sites where PG growth is completed, a region where PG synthesis completes first in Phase I will become a future cell pole.

### The MglA-MglB polarity axis regulates the distribution of the Rod system

As the Rod complex is the major system for PG growth in Phase II, the MglA-MglB polarity axis might regulate cell elongation through the Rod complexes. However, MglA and MglB are both cytoplasmic proteins, which are not likely to regulate the periplasmic activities of the Rod system directly. To visualize the distribution of the Rod complexes, we fused a DNA sequence encoding mCherry to the endogenous *rodA* gene in the wild-type background. The resulted strain showed minor delay of elongation during sporulation and was able to establish rod shape, indicating that the mCherry-labeled RodA retains its enzymatic activity (*SI Appendix,* Fig. S2, Table S1). RodA-mCherry formed clusters during both vegetative growth and germination (Fig. 4A). When imaged at 3.33 Hz, RodA clusters showed typical diffusion in vegetative cells, with *D_RodA_* = (3.07 ± 1.97) × 10^-2^ μm^2^/s (n = 160) (*SI Appendix,* Movie. S7). Importantly, mecillinam treatment did not affect the diffusivity of RodA clusters ((3.35 ± 2.14) × 10^-2^ μm^2^/s, n = 58, *p* = 0.39) (*SI Appendix,* Movie. S8), consistent with a previous report in *E. coli* that the diffusion of PBP2 does not correlate with its catalytic activity (29). Despite its negligible effect on diffusion, mecillinam significantly inhibited the formation of RodA clusters. During 150-s of imaging at 0.33 Hz, untreated vegetative cells in the exponential phase formed 10.10 ± 3.41 RodA clusters/cell (n = 50). In the presence of mecillinam, this number decreased to 3.06 ± 1.06 clusters/cell (n = 71) (Fig. 4A). Based on the above observations, we reasoned that the formation of RodA clusters, rather than their diffusion, correlates with PG growth by the Rod complexes.

**Fig. 4.**
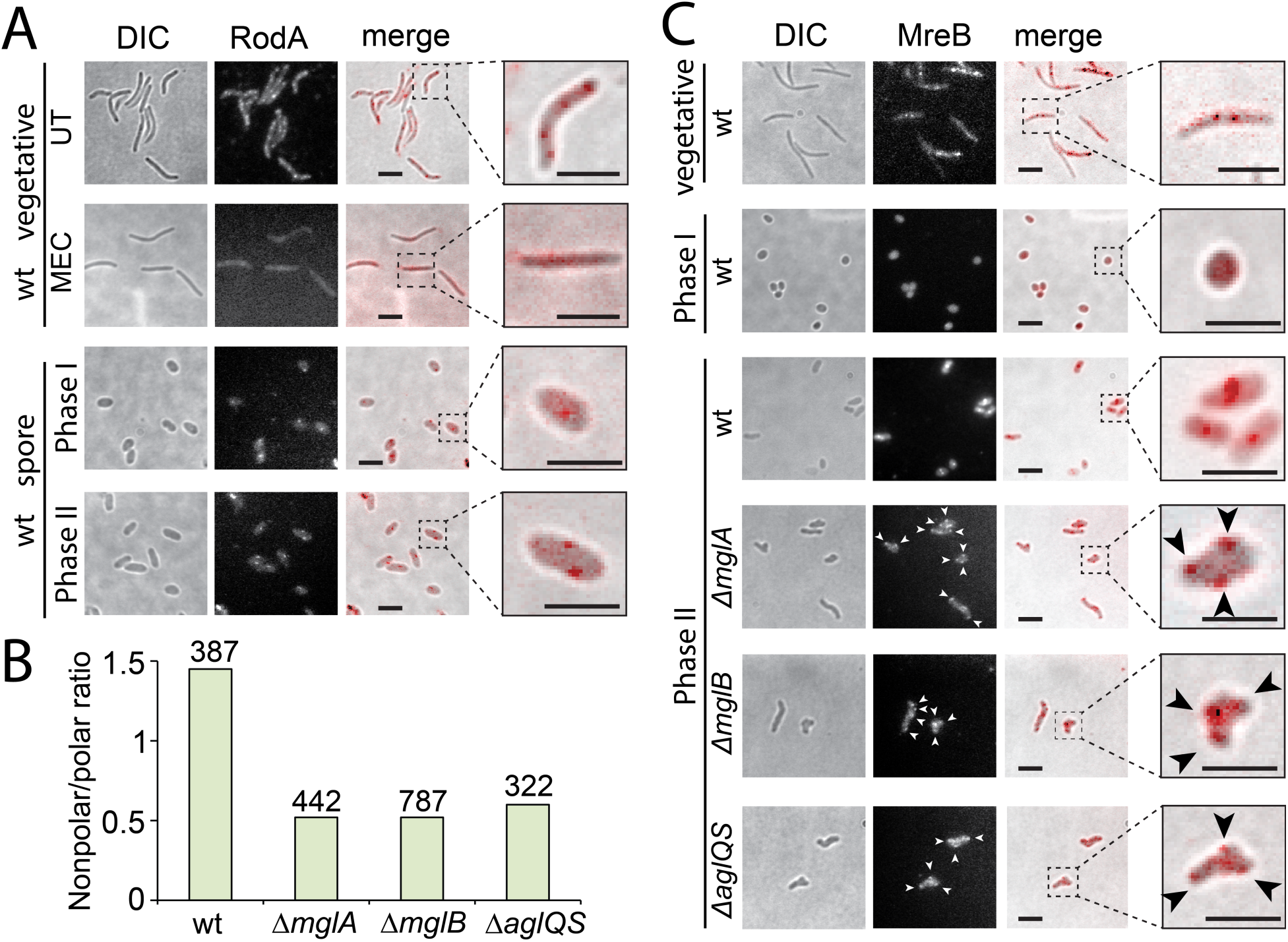
The MglA-MglB polarity axis regulates the distribution of Rod complexes through MreB and the gliding motor. **A)** RodA forms clusters during both vegetative growth and germination. Mecillinam reduces the formation of RodA clusters in vegetative cells. **B)** Nonpolar-to-polar distribution ratios of RodA clusters in the elongating cells of germination Phase II. The total number of RodA clusters analyzed is shown on top of each bar. **C)** The localization patterns of MreB filaments. Consistent with the altered distribution of RodA cluster, MreB patches (arrows) are frequently detected near cell poles and bulges in the emerging *ΔmglA*, *ΔmglB* and *ΔaglQS* cells in Phase II of germination. Scales bars, 2 μm

To investigate whether MglA and MglB regulate the distribution of the Rod complexes in Phase II of germination, we expressed RodA-mCherry endogenously in *ΔmglA* and *ΔmglB* spores. Along the long axis of the emerging cells, we loosely defined a region within 320 nm from each end of cell as a “pole” (which contains the cell pole and its adjacent subpolar region) and the rest of the cell as the “nonpolar region”. In the emerging cells from wild-type spores that expressed RodA-mCherry, the ratio between nonpolar and polar-localized RodA clusters was 1.45 (n = 387, Fig. 4B). In contrast, in the *ΔmglA* and *ΔmglB* backgrounds, this ratio decreased to 0.52 (n = 442) and 0.53 (n = 787), respectively (Fig. 4B). Our data support the hypothesis that during the sphere-to-rod transition, the established MglA-MglB axis expels the Rod system from cell poles.

### The MglA-MglB polarity axis regulates the distribution of Rod complexes through MreB and the gliding motor

The diffusion of RodA and MglB clusters is not likely connected. First, *D_RodA_* is two orders of magnitude higher than *D_MglB_*. Second, MglB only diffuses in Phase I of germination while RodA diffuses during both germination and vegetative growth. Thus, the MglA-MglB polarity axis does not likely regulate the distribution of Rod complexes directly. Since MglA and the Rod complex both bind to MreB, we hypothesized that the MglA-MglB polarity axis could regulate the distribution of Rod complexes through MreB. Although a photoactivatable mCherry (PAmCherry)-labeled MreB variant fully supports wild-type growth rate in vegetative cells (11), when expressed as the sole source of MreB, it failed to support rapid cell elongation in Phase II of germination. Nevertheless, spores expressing MreB-PAmCherry as merodiploids showed wild-type germination kinetics (*SI Appendix,* Fig. S2, Table S1), which were used to visualize the localization of MreB. When exposed to 405-nm excitation (0.2 kW/cm^2^) for 2 s, the majority of PAmCherry was photoactivated (11, 13). MreB-PAmCherry localized diffusively in Phase I spores, excluding the possibility that localized MreB filaments predetermine the polarity of spores. MreB started to form small patches in Phase II (Fig. 4C). Compared to the wild-type spores where MreB patches mainly localized at nonpolar locations in germination Phase II, many MreB patches formed near cell poles and bulges of the emerging *ΔmglA* and *ΔmglB* cells (Fig. 4C), consistent with the altered distribution of RodA clusters. While MreB might localize at bulges in response to altered local cell curvatures in these mutants (30–35), the aggregation of MreB at their poles is more likely due to the loss of the MglA-MglB polarity axis, because the poles of both wild-type and mutant cells have similar curvatures.

MglA connects MreB to the gliding motors and the gliding motors drive the movement of MreB filaments (5, 7, 11). To test if MglA recruits the gliding motors to transport the Rod complexes to nonpolar locations through MreB, we investigated the regrowth process of the *ΔaglQS* pseudospores that are sonication-sensitive due to the lack compact polysaccharide layers on their surfaces (36). *ΔaglQS* cells carry truncated gliding motors that are unable to drive the rapid motion of MreB filaments (11). Phenocopying the *ΔmglA* and *ΔmglB* spores, elongation of *ΔaglQS* pseudospores delayed significantly (Fig. 2A, 2B). Many emerging *ΔaglQS* cells displayed bulged surfaces in Phase II and survived at reduced rate, especially after hypoosmotic shock (Fig. 2A, 2E). Consistently, significantly higher PG growth was observed at cell poles and bulges in the elongation phase, similar to the observation made in *ΔmglA ΔdacB* and *ΔmglB ΔdacB* spores (Fig. 2D). Accordingly, significantly higher fractions of RodA clusters and MreB filaments were observed at poles and bulges in elongating *ΔaglQS* cells (Fig. 4B, 4C). In summary, MglA and MglB restrict PG growth to nonpolar regions in germination Phase II utilizing the gliding motors, which transport MreB, and thus the whole Rod complexes, under the control of MglA.

## Discussion

As spheres and rods are among the most common shapes adopted by walled bacteria, the sphere-to-rod transition during *M. xanthus* spore germination provides a unique opportunity to study rod-like morphogenesis in bacteria. Due to the absence of PG, glycerol-induced *M. xanthus* spores are especially valuable for the study of *de novo* PG synthesis, which drives spontaneous cell elongation in homogenous environments. We observed that MglB forms wandering clusters in Phase I of germination and that an emerging cell starts to elongate when the MglB cluster stops moving and stabilizes at what is to become a future pole. The dynamics of MglB clusters is sensitive to the status of PG synthesis because active PG growth prevents MglB clusters from settling down. Since MglB avoids colocalizing with MglA-GTP by converting the latter to MglA-GDP (4, 6), and MglA-GTP binds to MreB (which also carries the Rod complexes) (7), MglB clusters cannot stabilize at the sites where active PG synthesis by the Rod complex occurs. We propose that the wandering dynamics of MglB clusters serves as a mechanism to survey the status of PG growth and the region where PG growth completes first in Phase I will host a MglB cluster and become a future pole in Phase II (Fig. 5). Once an MglB cluster stabilizes at one pole, the expulsion between MglB and MglA-GTP causes MglA-GTP to cluster at the opposite side of the spore. At the poles that contain MglB clusters, MglB expels MglA-GTP and thus the Rod complexes from the poles. At the opposite poles, MglA-GTP stimulates the assembly of the gliding machinery by directly connecting it to MreB (7, 14). Once assembled, the gliding machineries transport MreB filaments, thus entire Rod complexes, away from the poles (5, 11) (Fig. 5). Taken together, the diametrically opposing clusters of MglA-GTP and MglB establish the polarity axis of the emerging cell, which restricts the activity of the Rod complex to nonpolar locations (Fig. 5). Other regulators that modulate the gliding motors, such as PlpA and RomR, could also connect to the Rod system through MreB, and thus respond to the status of PG synthesis. Utilizing the same system that defines the leading-lagging axis in vegetative cells, spherical spores grow their walls at nonpolar regions and eventually form rods.

**Fig.5.**
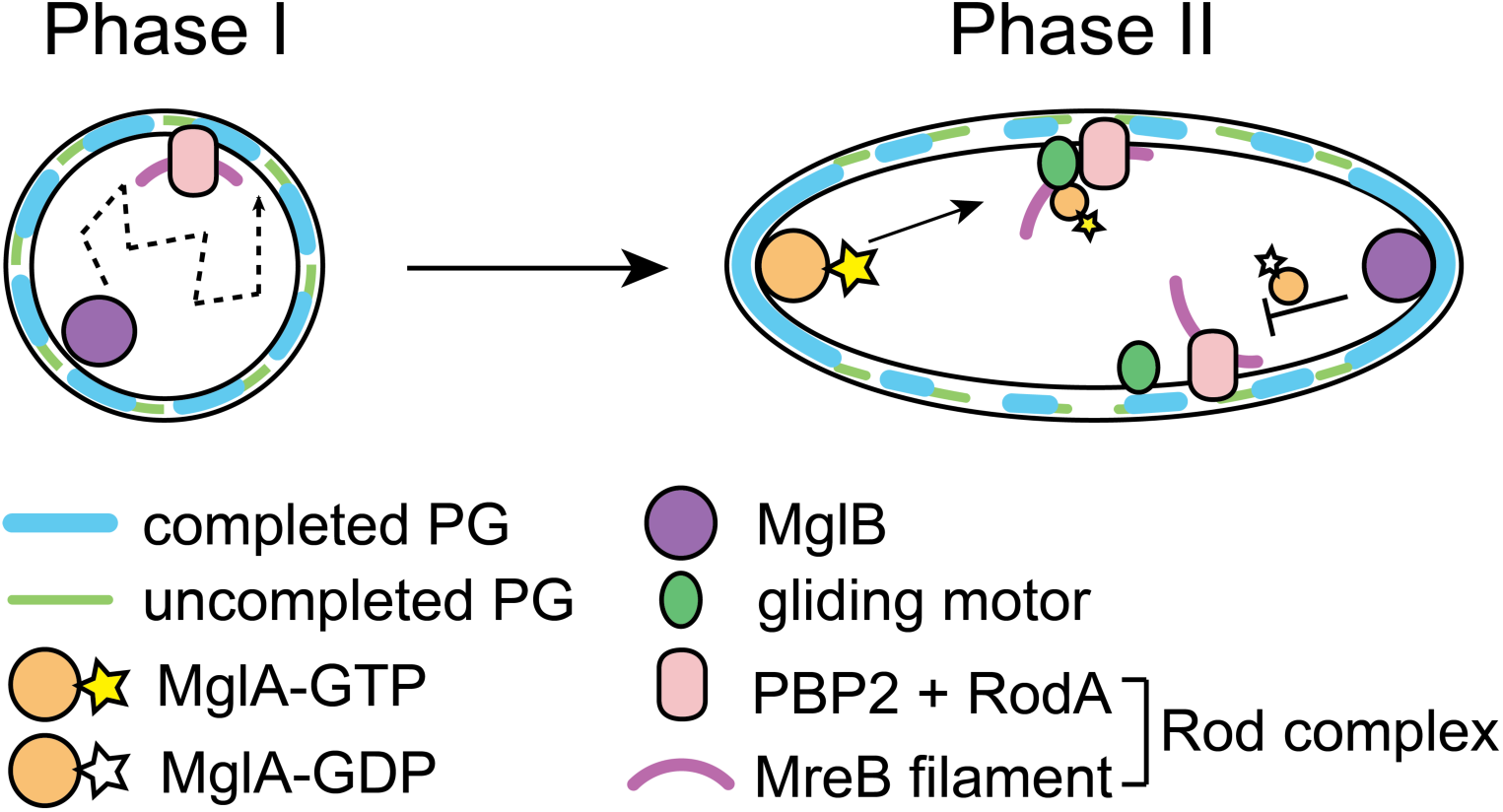
A schematic model for the *de novo* establishment of rod-shape by the MglA-MglB polarity axis.

Our findings suggest that multiple factors contribute to the establishment of rod shapes, among which the MglA-MglB polarity axis plays a prominent role. In the absence of MglA and MglB, other polarity regulators, including PlpA and RomR that also regulate the gliding motors, and potentially asymmetrically-localizes cellular components such as to-be-identified germinant receptors, ribosomes and RNA polymerase, still drive cell elongation, albeit much slower. Understanding these failsafe mechanisms will pave the way for fully understanding the mechanism of rod-like morphogenesis. The mechanism by which the gliding machineries facilitate both gliding and PG growth remains to be investigated. Nonetheless, our results have added another layer to the striking versatility of the gliding motors, which transport various cargos in different compartments of the cells: spore coats on cell surfaces (36), the Rod complex and some gliding proteins in the membrane and periplasm (12, 14), as well as MreB and other gliding proteins in the cytoplasm (11, 14, 37).

## Materials and Methods

### Strain construction

Deletion and insertion mutants were constructed by electroporating *M. xanthus* cells with 4 μg plasmid DNA or 1 μg of chromosomal DNA. Transformed cells were plated on CYE plates supplemented with 100 µg/ml sodium kanamycin sulfate or 10 µg/ml tetracycline hydrochloride. In-frame deletion of *dacB* is described in *SI Appendix, Materials and Methods, Table S2*.

### Sporulation, spore purification and germination

Vegetative *M. xanthus* cells were grown in liquid CYE medium (10 mM MOPS pH 7.6, 1% (w/v) Bacto™ casitone (BD Biosciences), 0.5% yeast extract and 4 mM MgSO_4_) at 32 °C, in 125-ml flasks with rigorous shaking, or on CYE plates that contains 1.5% agar. When liquid cell culture reaches OD_600_ 0.1 – 0.2, glycerol was added to 1 M to induce sporulation. After rigorous shaking overnight at 32 °C, remaining vegetative cells were eliminated by sonication and sonication-resistant spores were purified by centrifugation (1 min, 15,000 g and 4 °C). The pellet was washed three times with water. More details of spore purification and the purification of *ΔaglQS* pseudospores are provided in *SI Appendix, Materials and Methods*.

### Microscopy Analysis

Cryo-ET was performed on a Polara G2™ electron microscope. Images were collected at 9,000× magnification and 8-μm defocus, resulting in 0.42 nm/pixel. Data were acquired automatically with the SerialEM software (38). Time-lapse videos of the germination progress of wild-type and *ΔmglA* spores were recorded using an OMAX™ A3590U CCD camera and a Plan Flour™ 40×/0.75 Ph2 DLL objective on a phase-contrast Nikon Eclipse™ 600 microscope. The length, width and geometric aspect ratios (L/W) of spores/cells were determined from differential interference contrast (DIC) images using a custom algorithm written in MATLAB (The MathWorks, Inc., Natick, MA), which is available upon request. DIC images of spores/cells were captured using a Hamamatsu ImagEM X2™ EM-CCD camera C9100-23B (effective pixel size 160 nm) on an inverted Nikon Eclipse-Ti™ microscope with a 100× 1.49 NA TIRF objective, which are also used for capturing fluorescence images. MglB and RodA clusters were localized using an algorithm written in MATLAB (11), which is available upon request. More detailed information is provided in *SI Appendix, Materials and Methods*.

## ACKNOWLEDGEMENTS

We thank Autumn Ridge and Elias Topo for technical assistance, Drs. Joseph Sorg and Ritu Shrestha for their help in the initial determination of germination phenotypes, Drs. Michael Van Nieuwenhze and Yen-Pang Hsu for providing TADA, and Drs. David Zusman, Michael Manson and Joseph Sorg for critical reading of this manuscript. This work is supported by the National Institute of Health R01GM129000 (to B.N.) and R01AI087946 (to J.L.) and by CNRS to T.M.

## Notes

#### Summary of Updates

New experimental results presented Introduction and discussion revised Supplemental files updated

## References

1. S. van Teeffelen, L. D. Renner, Recent advances in understanding how rod-like bacteria stably maintain their cell shapes. F1000Res 7, 241 (2018).

2. H. Cho et al., Bacterial cell wall biogenesis is mediated by SEDS and PBP polymerase families functioning semi-autonomously. Nat Microbiol 10.1038/nmicrobiol.2016.172, 16172 (2016).

3. T. K. Lee, K. Meng, H. Shi, K. C. Huang, Single-molecule imaging reveals modulation of cell wall synthesis dynamics in live bacterial cells. Nature communications 7, 13170 (2016).

4. S. Leonardy et al., Regulation of dynamic polarity switching in bacteria by a Ras-like G-protein and its cognate GAP. Embo J 29, 2276–2289 (2010).

5. B. Nan et al., The polarity of myxobacterial gliding is regulated by direct interactions between the gliding motors and the Ras homolog MglA. Proc Natl Acad Sci U S A 112, E186–193 (2015).

6. Y. Zhang, M. Franco, A. Ducret, T. Mignot, A bacterial Ras-like small GTP-binding protein and its cognate GAP establish a dynamic spatial polarity axis to control directed motility. PLoS Biol 8, e1000430 (2010).

7. A. Treuner-Lange et al., The small G-protein MglA connects to the MreB actin cytoskeleton at bacterial focal adhesions. J Cell Biol 210, 243–256 (2015).

8. B. Nan, M. J. McBride, J. Chen, D. R. Zusman, G. Oster, Bacteria that glide with helical tracks. Curr Biol 24, R169–R173 (2014).

9. E. M. Mauriello et al., Bacterial motility complexes require the actin-like protein, MreB and the Ras homologue, MglA. Embo J 29, 315–326 (2010).

10. B. Nan, Bacterial Gliding Motility: Rolling Out a Consensus Model. Curr Biol 27, R154–R156 (2017).

11. G. Fu et al., MotAB-like machinery drives the movement of MreB filaments during bacterial gliding motility. Proc Natl Acad Sci U S A 115, 2484–2489 (2018).

12. B. Nan et al., Myxobacteria gliding motility requires cytoskeleton rotation powered by proton motive force. Proc Natl Acad Sci U S A 108, 2498–2503 (2011).

13. B. Nan et al., Flagella stator homologs function as motors for myxobacterial gliding motility by moving in helical trajectories. Proc Natl Acad Sci U S A 110, E1508–1513 (2013).

14. L. M. Faure et al., The mechanism of force transmission at bacterial focal adhesion complexes. Nature 539, 530–535 (2016).

15. D. R. Zusman, A. E. Scott, Z. Yang, J. R. Kirby, Chemosensory pathways, motility and development in *Myxococcus xanthus*. Nat Rev Microbiol 5, 862–872 (2007).

16. M. Dworkin, S. M. Gibson, A System for Studying Microbial Morphogenesis: Rapid Formation of Microcysts in *Myxococcus xanthus*. Science 146, 243–244 (1964).

17. E. I. Tocheva et al., Peptidoglycan transformations during *Bacillus subtilis* sporulation. Mol Microbiol 88, 673–686 (2013).

18. K. Khanna et al., The molecular architecture of engulfment during *Bacillus subtilis* sporulation. Elife 8 (2019).

19. N. K. Bui et al., The peptidoglycan sacculus of *Myxococcus xanthus* has unusual structural features and is degraded during glycerol-induced myxospore development. J Bacteriol 191, 494–505 (2009).

20. A. Vigouroux et al., Class-A penicillin binding proteins do not contribute to cell shape but repair cell-wall defects. Elife 9 (2020).

21. E. Kuru et al., Mechanisms of incorporation for D-amino acid probes that target peptidoglycan biosynthesis. ACS Chem Biol 10.1021/acschembio.9b00664 (2019).

22. Y. P. Hsu et al., Full color palette of fluorescent d-amino acids for in situ labeling of bacterial cell walls. Chem Sci 8, 6313–6321 (2017).

23. E. Kuru et al., In Situ probing of newly synthesized peptidoglycan in live bacteria with fluorescent D-amino acids. Angew Chem Int Ed Engl 51, 12519–12523 (2012).

24. M. F. Dion et al., Bacillus subtilis cell diameter is determined by the opposing actions of two distinct cell wall synthetic systems. Nat Microbiol 10.1038/s41564-019-0439-0 (2019).

25. D. Keilberg, K. Wuichet, F. Drescher, L. Sogaard-Andersen, A response regulator interfaces between the Frz chemosensory system and the MglA/MglB GTPase/GAP module to regulate polarity in *Myxococcus xanthus*. PLoS Genet 8, e1002951 (2012).

26. C. B. Pogue, T. Zhou, B. Nan, PlpA, a PilZ-like protein, regulates directed motility of the bacterium *Myxococcus xanthus*. Mol Microbiol 107, 214–228 (2018).

27. Y. Zhang, M. Guzzo, A. Ducret, Y. Z. Li, T. Mignot, A dynamic response regulator protein modulates G-protein-dependent polarity in the bacterium *Myxococcus xanthus*. PLoS Genet 8, e1002872 (2012).

28. T. Zhou, B. Nan, Exopolysaccharides promote *Myxococcus xanthus* social motility by inhibiting cellular reversals. Mol Microbiol 103, 729–743 (2017).

29. T. K. Lee et al., A dynamically assembled cell wall synthesis machinery buffers cell growth. Proc Natl Acad Sci U S A 111, 4554–4559 (2014).

30. B. P. Bratton, J. W. Shaevitz, Z. Gitai, R. M. Morgenstein, MreB polymers and curvature localization are enhanced by RodZ and predict *E. coli*’s cylindrical uniformity. Nature communications 9, 2797 (2018).

31. A. Colavin, H. Shi, K. C. Huang, RodZ modulates geometric localization of the bacterial actin MreB to regulate cell shape. Nature communications 9, 1280 (2018).

32. S. Hussain et al., MreB filaments align along greatest principal membrane curvature to orient cell wall synthesis. Elife 7 (2018).

33. T. S. Ursell et al., Rod-like bacterial shape is maintained by feedback between cell curvature and cytoskeletal localization. Proc Natl Acad Sci U S A 111, E1025–1034 (2014).

34. F. Wong, E. C. Garner, A. Amir, Mechanics and dynamics of translocating MreB filaments on curved membranes. Elife 8 (2019).

35. R. M. Morgenstein et al., RodZ links MreB to cell wall synthesis to mediate MreB rotation and robust morphogenesis. Proc Natl Acad Sci U S A 10.1073/pnas.1509610112 (2015).

36. M. Wartel et al., A versatile class of cell surface directional motors gives rise to gliding motility and sporulation in *Myxococcus xanthus*. PLoS Biol 11, e1001728 (2013).

37. M. Sun, M. Wartel, E. Cascales, J. W. Shaevitz, T. Mignot, Motor-driven intracellular transport powers bacterial gliding motility. Proc Natl Acad Sci U S A 108, 7559–7564 (2011).

38. D. N. Mastronarde, Automated electron microscope tomography using robust prediction of specimen movements. Journal of Structural Biology 152, 36–51 (2005).

