## supplementary text and figures for "Establishing Rod-Shape from Spherical, Peptidoglycan-Deficient Bacterial Spores"

### ***SI APPENDIX***

#### MATERIALS AND METHODS

##### Strain Construction

In-frame deletion cassettes of *dacB* and the insertion cassette for RodA-labeling were amplified with polymerase chain reaction (PCR) using chromosomal DNA as template, digested and inserted into plasmid pBJ113 (1) to produce pBN$_{\Delta dacB}$ and pBN-*rodA-mCherry*. For the labeling of RodA, Protein sequence SGGGGSGGGGSGGGGS was used as the linker between RodA and mCherry. All constructs were confirmed by DNA sequencing. Transformants were obtained by homologous recombination and confirmed by PCR. Strains, plasmids and PCR primers are listed in Table S2.

##### Immunoblot analysis

For each strain, cells were grown in CYE medium to $OD_{600}$ 1.0. 20-μl culture were lysed using 2× SDS loading buffer, subjected to electrophoresis using 4 – 15% gradient gels (Bio-rad) and blotted onto Amersham™ Hybond™ 0.2 μm PVDF blotting membranes (GE healthcare). MglB and MglB-mCherry were detected using anti-MglB antibodies (2); mCherry and RodA-mCherry, monoclonal anti-mCherry antibodies (abcam). Protein bands were visualized using horseradish peroxidase-conjugated goat-anti-rabbit secondary antibodies (Thermo Sceintific), the Pierce™ ECL blotting substrate (Thermo Sceintific), and the Amersham Hyperfilm™ ECL chemiilluminescence films (GE Healthcare).

##### Sporulation, spore purification and germination

To eliminate vegetative cells from spores, 400 μl of cell culture was transferred to a 1.5-ml microcentrifuge tube, sonicated eight times on ice for 2 s each, at 2-s intervals. The elimination of vegetative cells was confirmed by DIC or phase contrast microscopy. Additional sonication cycles were applied when vegetative cells still remained.

The *ΔaglQS* pseudospores are nonresistant to sonication (3). Nevertheless, as we used the aspect ratio L/W, rather than OD, to quantify symmetry-breaking, the observed delay in morphological transition is not likely due to the low survival rate of *ΔaglQS* pseudospores. In addition, these surviving *ΔaglQS* pseudospores regrew into rods in the same two-phase manner as the wild-type spores (Fig. 2). The *ΔaglQS* pseudospores were purified by centrifugation with sucrose gradient. 1 ml culture that contains *ΔaglQS* pseudospores was first collected by centrifugation (20 min, 1,800 g and room temperature). Then the pellet was washed three times with water, suspended in 2 ml water, pipetted to the top layer of a 35-ml centrifuge tube that contained 8 ml of 60% sucrose solution, and sedimented by centrifugation (20 min, 1,800 g and room temperature). The pellet was collected and washed with water five times.

Purified spores and pseudospores were suspended in water. Their ODs were measured at 600 nm and diluted to 0.5. To induce germination and regrowth, 1 ml of spores/pseudospores were collected again by centrifugation (1 min, 15,000 g and room temperature), suspended in 1 ml of liquid germination CYE (CYE medium supplemented with additional 0.2% casitone and 1 mM $CaCl_2$) and incubated in an 18-mm test tube at 32 °C, with vigorous shaking. To measure the germination and regrowth rates of the *pilA::tet*, *ΔmglA pilA::tet*, *ΔmglB pilA::tet* spores and the *ΔaglQS pilA::tet* pseudospores, spores/pseudospores were enumerated in bacterial cell counting chambers and dilution plated on solid CYE agar. Colonies were then counted after 120-h incubation at 32 °C. Hypoosmotic shock was performed on Phase II spores/pseudospores. Spores/pseudospores were enumerated, suspended in 1 ml of liquid germination CYE and incubated at 32 °C for 1 h, with vigorous shaking. Germinating spores/pseudospores were washed three times using 20 mM Hris-HCl pH7.6 and incubated in the same buffer for 1 h before being plated on CYE agar. Colonies were then counted after 120-h incubation at 32 °C. For inhibitor treatments, the minimum inhibitory concentration (MIC) of each inhibitor was determined on CYE agar using wild-type vegetative *M. xanthus* cells. For each inhibitor, 2 × MIC was used in both germination assay and TADA-labeling.

**Bright field microscopy and cell geometry analysis**

5 μl of spore/cell suspension was spotted on a germination CYE agar (1.5%) pad of ~0.5-mm thickness. Time-lapse videos of the germination progress of wild-type and *ΔmglA* spores were recorded using an OMAX™ A3590U CCD camera and a Plan Flour™ 40×/0.75 Ph2 DLL objective on a phase-contrast Nikon Eclipse™ 600 microscope. Germination temperature was maintained at 32 °C using an AmScope™ TCS-100 slide warmer. The length, width and geometric aspect ratios (L/W) of spores/cells were determined from differential interference contrast (DIC) images using a custom algorithm written in MATLAB (The MathWorks, Inc., Natick, MA), which is available upon request. DIC images of spores/cells were captured using a Hamamatsu ImagEM X2™ EM-CCD camera C9100-23B (effective pixel size 160 nm) on an inverted Nikon Eclipse-Ti™ microscope with a 100× 1.49 NA TIRF objective.

**Cryo-ET**

Vegetative cells and glycerol-induced spores of wild-type *M. xanthus* were mixed with BSA-golds as the fiducial marker before being transferred onto EM grids. The samples on EM grids were blotted by Whatman filter paper and rapidly plunge-frozen in liquid ethane in a homemade plunger apparatus (4). The hydrated samples on EM grids were transferred into liquid nitrogen before imaging. EM grids were then transferred to a Polara G2™ electron microscope. Images were collected at 9,000× magnification and 8-μm defocus, resulting in 0.42 nm/pixel. Data were acquired automatically with the SerialEM software (5). A total dose of 50 e/Å$^2$ was distributed among 35 tilt images covering angles from -51° to 51° at tilt steps of 3°. For every single tilt series collection, the dose-fractionated mode was used to generate 8 to 10 frames per projection image. Collected dose-fractionated data were first subjected to the motion correction program to generate drift-corrected stack files (6, 7). Contrast transfer-function correction of individual tilt images was performed using the function of *ctfphaseflip* implemented in IMOD (8). Tilt series were aligned in IMOD using gold fiducial markers and the alignment stacks were binned at 2 times (0.82 nm/pixel) to generate tomograms using SIRT reconstruction (9, 10).

**TADA labeling**

Lyophilized TADA was dissolved in DMSO at 150 mM and stored at −20 °C. The labeling was performed at 32 °C, using 1 μl of the TADA solution for 1ml of germinating spores. To visualize PG growth in Phase I, we added TADA to the medium at the beginning of germination and allowed *ΔdacB* spores to germinate for 1 h. To visualize PG growth in Phase II, we allowed spores to germinate for 1 h before adding TADA into the medium, then imaged the pattern of PG growth after 1 h of incubation in the presence of TADA. To determine the enzymatic systems for PG growth during germination, we added the inhibitors of different PBPs together with TADA. The sample was then washed four times and resuspended with TPM buffer (10 mM Tris-HCl pH 7.6, 1 mM $KH_2PO_4$, 8 mM $MgSO_4$) before being transferred to a 0.8% agarose pad of ~0.5 mm thickness, which was prepared by heat-dissolving agarose in 10 mM MOPS pH 7.6. Imaging was completed within 30 min. To quantify the incorporation of TADA, we defined the ends of spores on the longest axis as $0^{o}$ and $180^{o}$ and measured the fluorescence intensity of TADA in a circular region of 480-nm (3 pixels) diameter every $45^{o}$ around the spore envelope (Fig. 1E).

**Fluorescence microscopy and data analysis**

Fluorescence and PALM images were captured using a Hamamatsu ImagEM X2™ EM-CCD camera C9100-23B on an inverted Nikon Eclipse-Ti™ microscope with a 100× 1.49 NA TIRF objective. For all the imaging experiments, 5 μl of spores/cells at different germination time points were spotted on an agarose pad. For the treatments with inhibitors, inhibitors were added into both the spore/cell suspension and agarose pads.

YFP mCherry and TADA were activated by 488-nm, 561-nm and 532-nm lasers (0.2 kW/$cm^2$), respectively. The incorporation of TADA into PG was quantified by ImageJ. For each spore or emerging cell, the highest fluorescence intensity of TADA was normalized as 20 and the average fluorescence intensities were calculated from 20 spores/cells.

MglB and RodA clusters were localized using an algorithm written in MATLAB (11), which is available upon request. The MglB and RodA clusters that remained in focus for 4-12 frames (60 - 220 s) were subjected to analysis. Among these clusters, the ones that explored areas smaller than 160 nm ×160 nm in a time period of 220 s were considered as stationary. The diffusive clusters were fit by a symmetric 2D Gaussian function, whose center was assumed to be the cluster's position (12). Their diffusion coefficient (*D*) was determined from a linear fit to the first four points of the mean squared displacement (MSD) using formula MSD = $y^0 + 4D\Delta t$ (13). MreB-PAmCherry was activated using a 405-nm laser (0.3 - 3 W/cm$^2$, 1s) and imaged using a 561-nm laser (0.2 kW/cm$^2$, 0.1 s) under near total internal reflection illumination (14).

**SUPPLEMENTARY FIGURES**

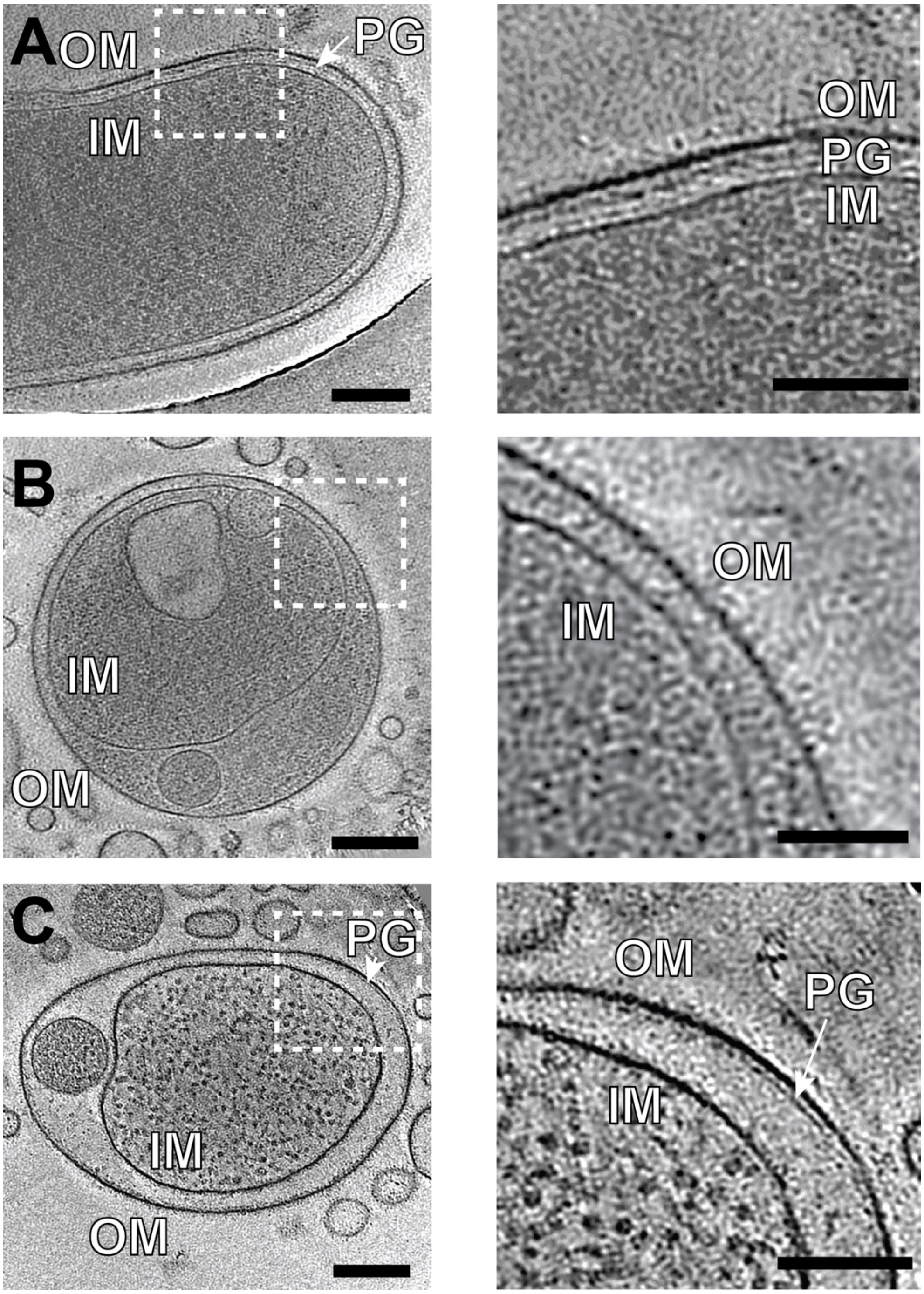


**Fig. S1. Mature *M. xanthus* spores do not retain intact PG layers that are sufficient to support cell shape.** Representative slices of 3D tomogram reconstructions of wild-type vegetative cells (**A**) and spores (**B**, **C**) are shown. While PG is clearly visible in vegetative cells (**A**), it is absent in glycerol-induced spores (**B**). **C)** Among 15 spores imaged, only one shows discontinuous densities that could represent PG fragments. Consistent with a pioneer study (15), vesicle-like structures were often observed between membranes, which could result from the excess membranes when rod-shaped cells convert to spherical spores (**B**, **C**). For each panel on the left (scale bars, 200 nm), a zoom-in view of the tomogram slice in a white dash box is shown on the right (scale bars, 50 nm).

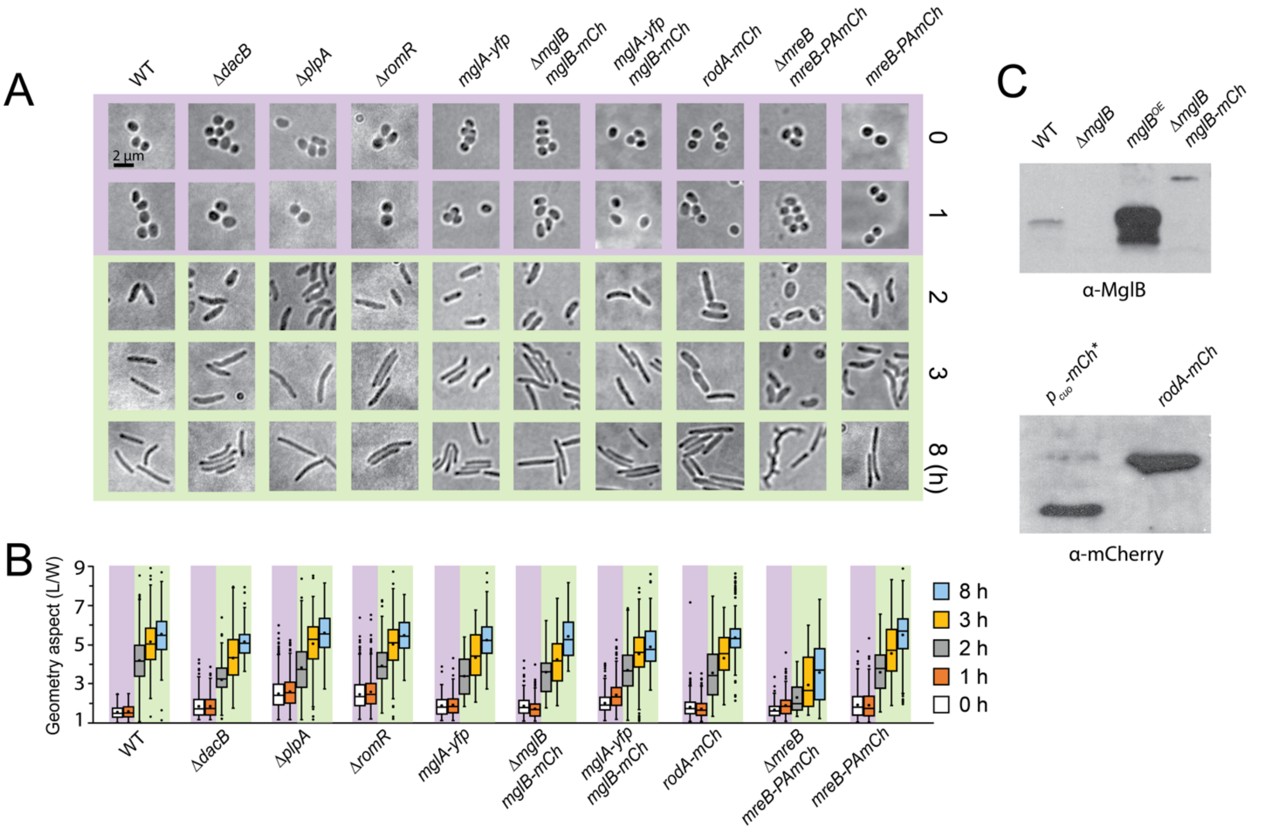


**Fig. S2. The germination of spores of various genetic backgrounds. A)** Morphological changes at different time points during germination. **B)** Quantitative analysis of the germination progress using the aspect ratios (L/W) of spores/cells. Boxes indicate the $25^{th}$ - $75^{th}$ percentiles, whiskers the $5^{th}$ - $95^{th}$ percentiles. In each box, the midline indicates the median and × indicates the mean (Table S1). Outlier data points are shown as individual dots above and below the whiskers. **C)** MglB-mCherry and RodA-mCherry are stably expressed in *M. xanthus* cells. *, leak expression.

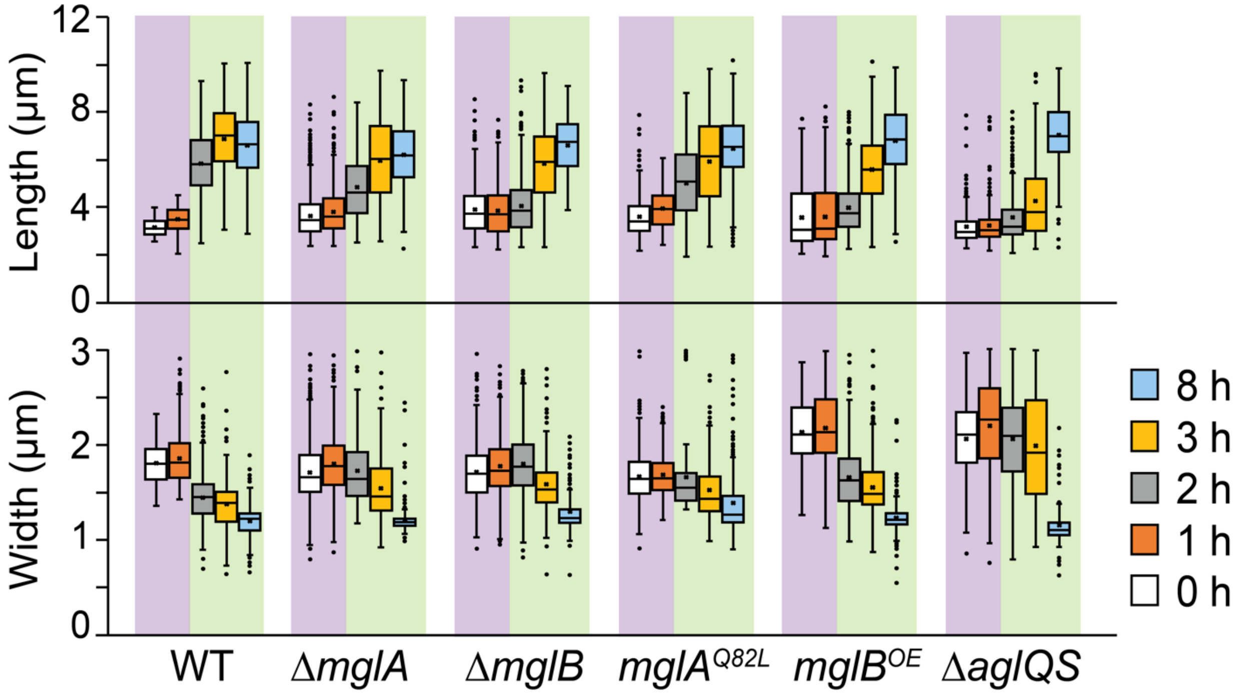


**Fig. S3. The changes of length and width during spore germination.** While the length and width of spores both increased slightly (by 6.91% and 2.74%, respectively) in Phase I of germination, maintaining the geometry aspect unchanged, when germination progressed to Phase II, emerging cells continued to grow in length but shrink in width.

#### MOVIE CAPTIONS

**Movie S1. The sphere-to-rod morphological transition during the germination of wild-type *M. xanthus* spores.** Images were taken at 4-min intervals and the movie plays at 10 Hz (2,400 × speedup).

**Movie S2. Oval spores do not preserve polarity from previous vegetative cells.** An oval spore elongates into rod shape along its short axis. Images were taken at 4-min intervals and the movie plays at 10 Hz (2,400 × speedup). Also see Fig. 1C.

**Movie S3. The *ΔmglA* spores generate pronounced bulges at nonpolar regions, appearing to have multiple cell poles.** Images were taken at 4-min intervals and the movie plays at 10 Hz (2,400 × speedup). Also see Fig. 2C.

**Movie S4. The diffusive dynamics of MglB-mCherry clusters in Phase I of germination.** Images were taken at 20-s intervals and the movie plays at 10 Hz (200 × speedup). Also see Fig. 3C.

**Movie S5. MglB-mCherry clusters stabilized at future cell poles during germination.** MglB-mCherry clusters first move randomly in spores then stabilize at future poles. Once MglB clusters stabilize, emerging cells start to elongate into rods. Images were taken at 5-min intervals and the movie plays at 5 Hz (1,500 × speedup). Also see Fig. 3D.

**Movie S6. MglB-mCherry clusters oscillate between cell poles in Phase II of germination.** Instead of diffusing, these MglB-mCherry clusters oscillate between cell poles. Images were taken at 20-s intervals and the movie plays at 10 Hz (200 × speedup). Also see Fig. 3C.

**Movie S7. The diffusion of RodA-mCherry clusters in untreated vegetative cells.** The diffusion of RodA-mCherry clusters Images were taken at 300-ms intervals and the movie plays at 10 Hz (3 × speedup).

**Movie S8. The diffusion of RodA-mCherry clusters in mecillinam-treated vegetative cells.** The diffusion of RodA-mCherry clusters Images were taken at 300-ms intervals and the movie plays at 10 Hz (3 × speedup).

**Table S1.** Quantification (mean ± SD) of germination progress of *M. xanthus* spores using their length (L, μm), width (W, μm), and length to width ratios (L/W) at different time points of germination.

| | | Germination time (h) | | | | |
|---|---|---|---|---|---|---|
| **Strain** | Treatment | **0** | **1** | **2** | **3** | **8** |
| Wild-type | Untreated | L/W = 1.56 ± 0.36<br>L = 3.33 ± 0.54<br>W = 1.82 ± 0.22<br>(n = 789[a])<br>[a], number of spores/cells | L/W = 1.55 ± 0.33<br>L = 3.56 ± 0.57<br>W = 1.87 ± 0.27<br>(n = 759) | L/W = 4.19 ± 1.21<br>L = 5.86 ± 1.32<br>W = 1.44 ± 0.27<br>(n = 412) | L/W = 5.16 ± 1.32<br>L = 6.88 ± 1.43<br>W = 1.38 ± 0.28<br>(n = 197) | L/W = 5.57 ± 1.08<br>L = 6.61 ± 1.41<br>W = 1.20 ± 0.18<br>(n = 232) |
| | Mecillinam (100 μg/ml) | | L/W = 1.48 ± 0.26<br>(n = 269) | L/W = 1.42 ± 0.28<br>(n = 261) | L/W = 1.45 ± 0.41<br>(n = 427) | L/W = 1.67 ± 0.47<br>(n = 213) |
| | A22 (10 μg/ml) | | L/W = 1.45 ± 0.37<br>(n = 696) | L/W = 1.57 ± 0.42<br>(n = 401) | L/W = 1.60 ± 0.45<br>(n = 443) | L/W = 1.50 ± 0.42<br>(n = 903) |
| | Cefsulodin (5 mg/ml) | | L/W = 1.57 ± 0.43<br>(n = 319) | L/W = 2.18 ± 0.68<br>(n = 221) | L/W = 2.95 ± 0.89<br>(n = 265) | L/W = 1.96 ± 0.99<br>(n = 339) |
| | Cefmetazole (5 mg/ml) | | L/W = 1.75 ± 0.57<br>(n = 141) | L/W = 2.08 ± 0.73<br>(n = 233) | L/W = 2.72 ± 1.18<br>(n = 562) | L/W = 1.19 ± 0.38<br>(n = 308) |
| | Fosfomycin (1 mg/ml) | | L/W = 1.57 ± 0.36<br>(n = 501) | L/W = 3.03 ± 1.00<br>(n = 446) | L/W = 3.86 ± 1.13<br>(n = 235) | L/W = 4.19 ± 1.21<br>(n = 252) |
| *mreB*$^{V323A}$ | A22 (10 μg/ml) | L/W = 2.40 ± 0.73<br>(n = 450) | L/W = 2.41 ± 0.78<br>(n = 576) | L/W = 2.97 ± 0.95<br>(n = 539) | L/W = 3.81 ± 1.08<br>(n = 608) | L/W = 4.70 ± 1.26<br>(n = 479) |
| *ΔmglA* | Untreated | L/W = 2.19 ± 0.69<br>L = 3.62 ± 0.86<br>W = 1.71 ± 0.29<br>(n = 2033) | L/W = 2.19 ± 0.73<br>L = 3.79 ± 0.92<br>W = 1.80 ± 0.31<br>(n = 1224) | L/W = 2.65 ± 1.05<br>L = 4.82 ± 1.34<br>W = 1.71 ± 0.33<br>(n = 471) | L/W = 4.09 ± 1.47<br>L = 5.93 ± 1.64<br>W = 1.54 ± 0.37<br>(n = 140) | L/W = 5.08 ± 1.02<br>L = 6.17 ± 1.31<br>W = 1.18 ± 0.07<br>(n = 194) |
| *ΔmglB* | Untreated | L/W = 1.97 ± 0.72<br>L = 3.90 ± 0.99<br>W = 1.70 ± 0.29<br>(n = 536) | L/W = 1.93 ± 0.77<br>L = 3.87 ± 1.00<br>W = 1.78 ± 0.32<br>(n = 511) | L/W = 2.02 ± 0.89<br>L = 4.06 ± 1.12<br>W = 1.80 ± 0.33<br>(n = 1181) | L/W = 3.84 ± 1.39<br>L = 5.82 ± 1.64<br>W = 1.59 ± 0.29<br>(n = 297) | L/W = 5.18 ± 1.01<br>L = 6.61 ± 1.16<br>W = 1.29 ± 0.21<br>(n = 161) |
| *mglA*$^{Q82L}$ | Untreated | L/W = 2.25 ± 0.68<br>L = 3.60 ± 0.82<br>W = 1.67 ± 0.27<br>(n = 383) | L/W = 2.27 ± 0.49<br>L = 3.95 ± 0.82<br>W = 1.68 ± 0.22<br>(n = 250) | L/W = 2.59 ± 0.92<br>L = 5.02 ± 1.53<br>W = 1.57 ± 0.22<br>(n = 212) | L/W = 4.10 ± 1.55<br>L = 5.94 ± 1.89<br>W = 1.52 ± 0.32<br>(n = 142) | L/W = 4.87 ± 1.32<br>L = 6.46 ± 1.37<br>W = 1.39 ± 0.34<br>(n = 273) |
| *mglB*$^{OE}$ | Untreated | L/W = 1.43 ± 0.37<br>L = 3.79 ± 1.25<br>W = 2.12 ± 0.33<br>(n = 352) | L/W = 1.53 ± 0.47<br>L = 3.81 ± 1.20<br>W = 2.16 ± 0.37<br>(n = 488) | L/W = 2.56 ± 0.82<br>L = 4.18 ± 1.53<br>W = 1.64 ± 0.33<br>(n = 572) | L/W = 3.82 ± 1.11<br>L = 5.72 ± 1.32<br>W = 1.54 ± 0.30<br>(n = 399) | L/W = 5.67 ± 1.18<br>L = 6.88 ± 1.52<br>W = 1.22 ± 0.19<br>(n = 202) |
| *ΔaglQS* | Untreated | L/W = 1.43 ± 0.45<br>L = 3.16 ± 0.76<br>W = 2.10 ± 0.38<br>(n = 259) | L/W = 1.38 ± 0.53<br>L = 3.21 ± 0.80<br>W = 2.23 ± 0.46<br>(n = 289) | L/W = 1.84 ± 1.03<br>L = 3.56 ± 1.14<br>W = 2.10 ± 0.45<br>(n = 264) | L/W = 2.35 ± 1.38<br>L = 4.25 ± 1.55<br>W = 2.03 ± 0.54<br>(n = 527) | L/W = 5.69 ± 1.24<br>L = 6.18 ± 1.20<br>W = 1.24 ± 0.21<br>(n = 131) |
| *ΔplpA* | Untreated | L/W = 2.29 ± 0.76<br>(n = 761) | L/W = 2.39 ± 0.75<br>(n = 784) | L/W = 3.56 ± 1.18<br>(n = 321) | L/W = 4.74 ± 1.37<br>(n = 124) | L/W = 5.27 ± 1.00<br>(n = 158) |
| *ΔromR* | Untreated | L/W = 2.33 ± 0.79<br>(n = 1238) | L/W = 2.46 ± 0.87<br>(n = 580) | L/W = 4.98 ± 1.64<br>(n = 216) | L/W = 4.99 ± 1.64<br>(n = 118) | L/W = 5.46 ± 1.03<br>(n = 162) |
| *ΔdacB* | Untreated | L/W = 1.63 ± 0.50<br>(n = 693) | L/W = 1.62 ± 0.47<br>(n = 390) | L/W = 2.84 ± 0.77<br>(n = 605) | L/W = 3.83 ± 1.27<br>(n = 350) | L/W = 4.57 ± 0.66<br>(n = 181) |
| *mglA-yfp* | Untreated | L/W = 1.79 ± 0.53<br>(n = 149) | L/W = 1.83 ± 0.55<br>(n = 271) | L/W = 3.36 ± 1.03<br>(n = 434) | L/W = 4.33 ± 1.25<br>(n = 577) | L/W = 5.25 ± 1.09<br>(n = 392) |
| *ΔmglB mglB-mCh* | Untreated | L/W = 1.99 ± 0.59<br>(n = 192) | L/W = 1.71 ± 0.50<br>(n = 575) | L/W = 3.83 ± 1.31<br>(n = 572) | L/W = 4.56 ± 1.60<br>(n = 614) | L/W = 5.89 ± 1.29<br>(n = 377) |
| *mglA-yfp mglB-mCh* | Untreated | L/W = 1.90 ± 0.53<br>(n = 348) | L/W = 2.31 ± 0.69<br>(n = 526) | L/W = 3.56 ± 1.12<br>(n = 501) | L/W = 4.39 ± 1.31<br>(n = 341) | L/W = 4.76 ± 1.05<br>(n = 472) |
| *rodA-mCh* | Untreated | L/W = 1.75 ± 0.53<br>(n = 477) | L/W = 1.66 ± 0.43<br>(n = 556) | L/W = 3.50 ± 1.31<br>(n = 467) | L/W = 4.25 ± 1.19<br>(n = 509) | L/W = 5.31 ± 1.16<br>(n = 423) |
| *ΔmreB mreB-PAmCh* | Untreated | L/W = 1.64 ± 0.43<br>(n = 131) | L/W = 1.87 ± 0.57<br>(n = 257) | L/W = 2.30 ± 0.86<br>(n = 564) | L/W = 2.96 ± 1.22<br>(n = 865) | L/W = 3.65 ± 1.75<br>(n = 566) |
| *mreB-PAmCh* | Untreated | L/W = 1.89 ± 0.67<br>(n = 252) | L/W = 1.89 ± 0.71<br>(n = 484) | L/W = 3.59 ± 1.16<br>(n = 281) | L/W = 4.56 ± 1.42<br>(n = 362) | L/W = 5.51 ± 1.38<br>(n = 274) |

**Table S2. Strains, primers and plasmids**

| Bacterial Strains | Source | Identifier |
|---|---|---|
| DZ2 (wild-type *M. xanthus* strain) | Laboratory stock | N/A |
| *pilA::tet* | (16) | DZ4469 |
| *mreB*$^{V323A}$ | (17) | TM264 |
| *ΔmglA* | (2) | TM12 |
| *ΔmglA pilA::tet* | (18) | BN220 |
| *ΔmglB* | (2) | TM155 |
| *ΔmglB pilA::tet* | (18) | BN221 |
| *ΔmglA* $p_{mglA}$-*mglA*$^{Q82L}$ | (2) | TM239 |
| *ΔplpA* | (18) | BN201 |
| *ΔromR* | (19) | TM254 |
| *ΔaglQS* | (18) | BN121 |
| *ΔaglQS pilA::tet* | This study | BN285 |
| *ΔdacB* | This study | TM1142 |
| *ΔmglA ΔdacB* | This study | BN286 |
| *ΔmglB ΔdacB* | This study | BN287 |
| *ΔaglQS ΔdacB* | This study | BN288 |
| *mglA-yfp* | (2) | TM17 |
| *ΔmglB mglA-yfp* | (2) | TM192 |
| *ΔmglB* $p_{mglB}$-*mglB-mCherry* | This study | BN289 |
| *mglA-yfp mglB-mCherry* | This study | BN290 |
| *ΔmreB* $p_{mreB}$-*mreB-PAmCherry* | (11) | BN291 |
| $p_{mreB}$-*mreB-PAmCherry* | This study | BN292 |
| *ΔmglA* $p_{mreB}$-*mreB-PAmCherry* | This study | BN293 |
| *ΔmglB* $p_{mreB}$-*mreB-PAmCherry* | This study | BN294 |
| *ΔaglQS* $p_{mreB}$-*mreB-PAmCherry* | This study | BN295 |
| *rodA-mCherry::kan* | This study | BN296 |
| $p_{cuo}$-*mCherry* | This study | BN297 |
| *ΔmglA rodA-mCherry::kan* | This study | BN298 |
| *ΔmglB rodA-mCherry::kan* | This study | BN299 |
| *ΔaglQS rodA-mCherry::kan* | This study | BN300 |
| Primers for the construction of pBN$_{ΔdacB}$ | | |
| CGGAATTCAGGTCCATGCCAATCAGCTC | This study | N/A |
| GGGGTACCGACTGGCCGCCTGGAAAGG | This study | N/A |
| CGGGATCCAGGACGCACCGTCTCATTC | This study | N/A |
| GCTCTAGAGGCCACGTCATAGACGGTG | This study | N/A |
| Primers for labeling RodA with mCherry | | |
| AAGCTTCGAGCGCGACCACGCCTGGTA | This study | N/A |
| GGATCCGAACATGTGACGGCGCATGCTGA | This study | N/A |
| Plasmids | | |
| pBJ113 | (1) | pBJ113 |
| pCK126 (pSWU30*mglB-mCherry*) | (20) | N/A |
| pBN-*mCherry* | (21) | N/A |
| pBN$_{mreB}$ *(pSWU30-*$P_{mreB}$*-mreB-PAmCherry)* | (11) | N/A |
| pBN$_{ΔdacB}$ (plasmid for *dacB* deletion) | This study | N/A |
| pBN-*rodA-mCherry* | This study | N/A |

#### References

1. B. Julien, A. D. Kaiser, A. Garza, Spatial control of cell differentiation in *Myxococcus xanthus*. *Proceedings of the National Academy of Sciences of the United States of America* **97**, 9098-9103 (2000).
2. Y. Zhang, M. Franco, A. Ducret, T. Mignot, A bacterial Ras-like small GTP-binding protein and its cognate GAP establish a dynamic spatial polarity axis to control directed motility. *PLoS Biol* **8**, e1000430 (2010).
3. M. Wartel *et al.*, A versatile class of cell surface directional motors gives rise to gliding motility and sporulation in *Myxococcus xanthus*. *PLoS Biol* **11**, e1001728 (2013).
4. S. Zhu, Z. Qin, J. Wang, D. R. Morado, J. Liu, In Situ Structural Analysis of the Spirochetal Flagellar Motor by Cryo-Electron Tomography. *Methods Mol Biol* **1593**, 229-242 (2017).
5. D. N. Mastronarde, Automated electron microscope tomography using robust prediction of specimen movements. *Journal of Structural Biology* **152**, 36-51 (2005).
6. X. M. Li *et al.*, Electron counting and beam-induced motion correction enable near-atomic-resolution single-particle cryo-EM. *Nature Methods* **10**, 584-+ (2013).
7. D. R. Morado, B. Hu, J. Liu, Using Tomoauto: A Protocol for High-throughput Automated Cryo-electron Tomography. *Jove-J Vis Exp* ARTN e53608 10.3791/53608 (2016).
8. Q. R. Xiong, M. K. Morphew, C. L. Schwartz, A. H. Hoenger, D. N. Mastronarde, CTF determination and correction for low dose tomographic tilt series. *Journal of Structural Biology* **168**, 378-387 (2009).
9. J. I. Agulleiro, J. J. Fernandez, Tomo3D 2.0-Exploitation of Advanced Vector eXtensions (AVX) for 3D reconstruction. *Journal of Structural Biology* **189**, 147-152 (2015).
10. J. R. Kremer, D. N. Mastronarde, J. R. McIntosh, Computer visualization of three-dimensional image data using IMOD. *Journal of Structural Biology* **116**, 71-76 (1996).
11. G. Fu *et al.*, MotAB-like machinery drives the movement of MreB filaments during bacterial gliding motility. *Proc Natl Acad Sci U S A* **115**, 2484-2489 (2018).
12. E. Betzig *et al.*, Imaging intracellular fluorescent proteins at nanometer resolution. *Science* **313**, 1642-1645 (2006).
13. T. K. Lee, K. Meng, H. Shi, K. C. Huang, Single-molecule imaging reveals modulation of cell wall synthesis dynamics in live bacterial cells. *Nature communications* **7**, 13170 (2016).
14. B. Nan *et al.*, Flagella stator homologs function as motors for myxobacterial gliding motility by moving in helical trajectories. *Proc Natl Acad Sci U S A* **110**, E1508-1513 (2013).
15. K. Bacon, F. A. Eiserling, A unique structure in microcysts of *Myxococcus xanthus*. *J Ultrastruct Res* **21**, 378-382 (1967).
16. H. C. Vlamakis, J. R. Kirby, D. R. Zusman, The Che4 pathway of *Myxococcus xanthus* regulates type IV pilus-mediated motility. *Mol Microbiol* **52**, 1799-1811 (2004).

17. E. M. Mauriello *et al.*, Bacterial motility complexes require the actin-like protein, MreB and the Ras homologue, MglA. *Embo J* **29**, 315-326 (2010).
18. C. B. Pogue, T. Zhou, B. Nan, PlpA, a PilZ-like protein, regulates directed motility of the bacterium *Myxococcus xanthus*. *Mol Microbiol* **107**, 214-228 (2018).
19. Y. Zhang, M. Guzzo, A. Ducret, Y. Z. Li, T. Mignot, A dynamic response regulator protein modulates G-protein-dependent polarity in the bacterium *Myxococcus xanthus*. *PLoS Genet* **8**, e1002872 (2012).
20. C. Kaimer, D. R. Zusman, Phosphorylation-dependent localization of the response regulator FrzZ signals cell reversals in *Myxococcus xanthus*. *Mol Microbiol* **88**, 740-753 (2013).
21. B. Nan, E. M. Mauriello, I. H. Sun, A. Wong, D. R. Zusman, A multi-protein complex from *Myxococcus xanthus* required for bacterial gliding motility. *Mol Microbiol* **76**, 1539-1554 (2010).
